# Millefy: visualizing cell-to-cell heterogeneity in read coverage of single-cell RNA sequencing datasets

**DOI:** 10.1101/537936

**Authors:** Haruka Ozaki, Tetsutaro Hayashi, Mana Umeda, Itoshi Nikaido

## Abstract

**Background:** Read coverage of RNA sequencing data reflects gene expression and RNA processing events. Single-cell RNA sequencing (scRNA-seq) methods, particularly “full-length” ones, provide read coverage of many individual cells and have the potential to reveal cellular heterogeneity in RNA transcription and processing. However, visualization tools suited to highlighting cell-to-cell heterogeneity in read coverage are still lacking.

**Results:** Here, we have developed Millefy, a tool for visualizing read coverage of scRNA-seq data in genomic contexts. Millefy is designed to show read coverage of all individual cells at once in genomic contexts and to highlight cell-to-cell heterogeneity in read coverage. By visualizing read coverage of all cells as a heat map and dynamically reordering cells based on diffusion maps, Millefy facilitates discovery of “local” region-specific, cell-to-cell heterogeneity in read coverage, including variability of transcribed regions.

**Conclusions:** Millefy simplifies the examination of cellular heterogeneity in RNA transcription and processing events using scRNA-seq data. Millefy is available as an R package (https://github.com/yuifu/millefy) and a Docker image to help use Millefy on the Jupyter notebook (https://hub.docker.com/r/yuifu/datascience-notebook-millefy).

## Background

Single-cell RNA sequencing (scRNA-seq) has been increasingly important in many areas, including developmental biology and cancer biology. In scRNA-seq data analyses, visualization is crucial for quality control (QC) as well as exploratory data analyses. For example, dimensionality reduction techniques such as principal component analysis (PCA) [1] or t-distributed stochastic neighbor embedding [2] are applied to a gene expression matrix to visualize individual cells as points in 2- or 3-dimensional space. Heat maps are also used to visualize gene expression matrices and highlight latent clusters of genes and cells. To date, many tools for visualizing gene expression matrices of scRNA-seq data have been proposed [3].

In contrast to visualization of gene expression matrices, visualization of read coverage, which is the distribution of mapped reads along genomic coordinates, helps reveal diverse aspects of RNA sequencing data and thus RNA biology and functional genomics. For example, read coverage reflects transcribed gene structures (e.g., exon-intron structures and transcript isoforms) [4], RNA processing events (e.g., normal and recursive splicing [5]), and transcription of intergenic and unannotated regions (e.g., enhancer RNAs [eRNAs]) [6]. Moreover, visual inspection of read coverage enables quality assessment of experimental methods (e.g., whether amplification is biased [7]) and bioinformatic methods (e.g., the accuracy of expression level estimation).

Given that scRNA-seq has revealed cellular heterogeneity in gene [8] and splicing isoform expression [9], [10], visualization of read coverage of scRNA-seq data is expected to reveal cellular heterogeneity in read coverage, which can be interpreted as biological (e.g., transcription and RNA processing) and technical (e.g., amplification biases) heterogeneity. Read coverage is informative, especially for so-called “full-length” scRNA-seq methods such as Smart-seq2 [11] and RamDA-seq [12], compared with “3′-tag sequencing” scRNA-seq methods, which sequence only the 3′ ends of RNAs and cannot be used to extract rich information from read coverage [13] [14]. Despite their potential importance, however, tools specifically for the visualization of read coverage of scRNA-seq data are still lacking.

To explore cell-to-cell heterogeneity in read coverage, we propose several requirements of a tool for visualization of read coverage in scRNA-seq data (Table 1). First, the tool must be able to display read coverage of all individual cells in a scRNA-seq dataset at once. This is because scRNA-seq data consist of many (10^2^−10^3^) cells and frequently includes latent heterogeneity that is masked by the summation of expression across cells. Second, the tool must associate read coverage with genomic contexts, such as gene structures and epigenomic features, because read coverage data can be interpreted only when it is displayed simultaneously with their genomic contexts. Third, the tool must highlight the cell-to-cell heterogeneity of read coverage within focal regions. This is because there should be “local” region-specific cell-to-cell heterogeneity in read coverage at transcriptional (e.g., antisense RNAs and eRNAs) and post-transcriptional (e.g., alternative splicing) levels, and such heterogeneity is difficult to notice in advance by cell groupings defined according to global similarity among cells.

**Table 1.** Comparison of Millefy to other visualization tools.

|  | Millefy | Genome browsers | Heatmaps |
| --- | --- | --- | --- |
| Visualize many cells at once | ✓ |  | ✓ |
| Associate read coverage with genomic contexts | ✓ | ✓ |  |
| Highlight cell-to-cell heterogeneity in read coverage | ✓ |  | ✓ |

Genome browsers and heat maps are two major tools for read coverage visualization. However, they are insufficient for fulfilling the above requirements.

Genome browsers, such as IGV [15] and JBrowse [16], utilize “tracks” to display various types of biological data, including gene annotations, positions of regulatory elements, and read mapping of next-generation sequencing (NGS) data along genomic coordinates. By stacking tracks in genome browsers, read coverage can easily be compared with other features and be interpreted in genomic contexts like gene models and epigenomic signals, which helps to generate and validate biological hypotheses. However, existing genome browsers are not suited for the large numbers of samples (i.e., cells) in scRNA-seq experiments. Indeed, efforts to visualize read coverage of scRNA-seq data using genome browsers have been limited to displays of a few dozen cells without the need to scroll [17] [18]. Although IGV and JBrowse implement heat map representations of tracks to show many cells at once, they cannot dynamically reorder tracks to reveal local cell-to-cell heterogeneity in read coverage.

Tools for heat maps combined with clustering algorithms have been used in the analysis of scRNA-seq data. Thus, heat maps can be used to visualize read coverage of all cells at once and reveal heterogeneity in read coverage. However, tools for generating heat maps are unsuited for visualizing read coverage of scRNA-seq data in genomic contexts, or they lack functionality to directly extract read coverage from standard NGS data formats.

Here, we have developed Millefy, which combines genome-browser-like visualization, heat maps, and dynamic reordering of single-cell read coverage and thus facilitates the examination of local heterogeneity within scRNA-seq data. Millefy extracts and organizes various types of useful information from read coverage of scRNA-seq data.

## Implementation

Millefy visualizes read coverage from each individual cell as a heat map in which rows represent cells and columns represent genomic bins within a focal region. The heat map is aligned with tracks for gene annotations, genomic features, and bulk NGS data, enabling comparisons of single-cell read coverage with genomic contexts. To highlight latent cell-to-cell heterogeneity in read coverage, the heat map rows (i.e., cells) are automatically and dynamically reordered by ‘local’ pseudo-time, which is calculated using diffusion maps [19], a nonlinear dimensionality reduction method. Specifically, diffusion maps are applied to matrixes of single-cell read coverage, where rows are cells and columns are genomic bins, and the first diffusion component is used to dynamically reorder cells either in an”all cells” manner or in a “group-wise” manner when groupings of cells are provided by users. Alternatively, PCA can also be used to reorder cells, but based on our experience, we recommend using diffusion maps (data not shown).

Millefy visualizations consist of five types of tracks (Figure 1): (1) scRNA-seq tracks, which display scRNA-seq read coverage as a heat map with ordered cells, (2) mean scRNA-seq read coverage tracks, (3) bulk NGS data tracks, which display read coverage of other NGS data, (4) BED tracks, which display genomic intervals defined by BED files, and (5) gene annotation tracks. In scRNA-seq tracks and bulk NGS data tracks, read coverage is normalized by user-provided normalization factors to correct for differences in the number of mapped reads among samples. Using the above tracks, Millefy can simultaneously display read coverage of each cell and mean read coverage of cells in each user-defined cell group as well as align scRNA-seq data with genome annotation data and NGS data.

**Figure 1.**
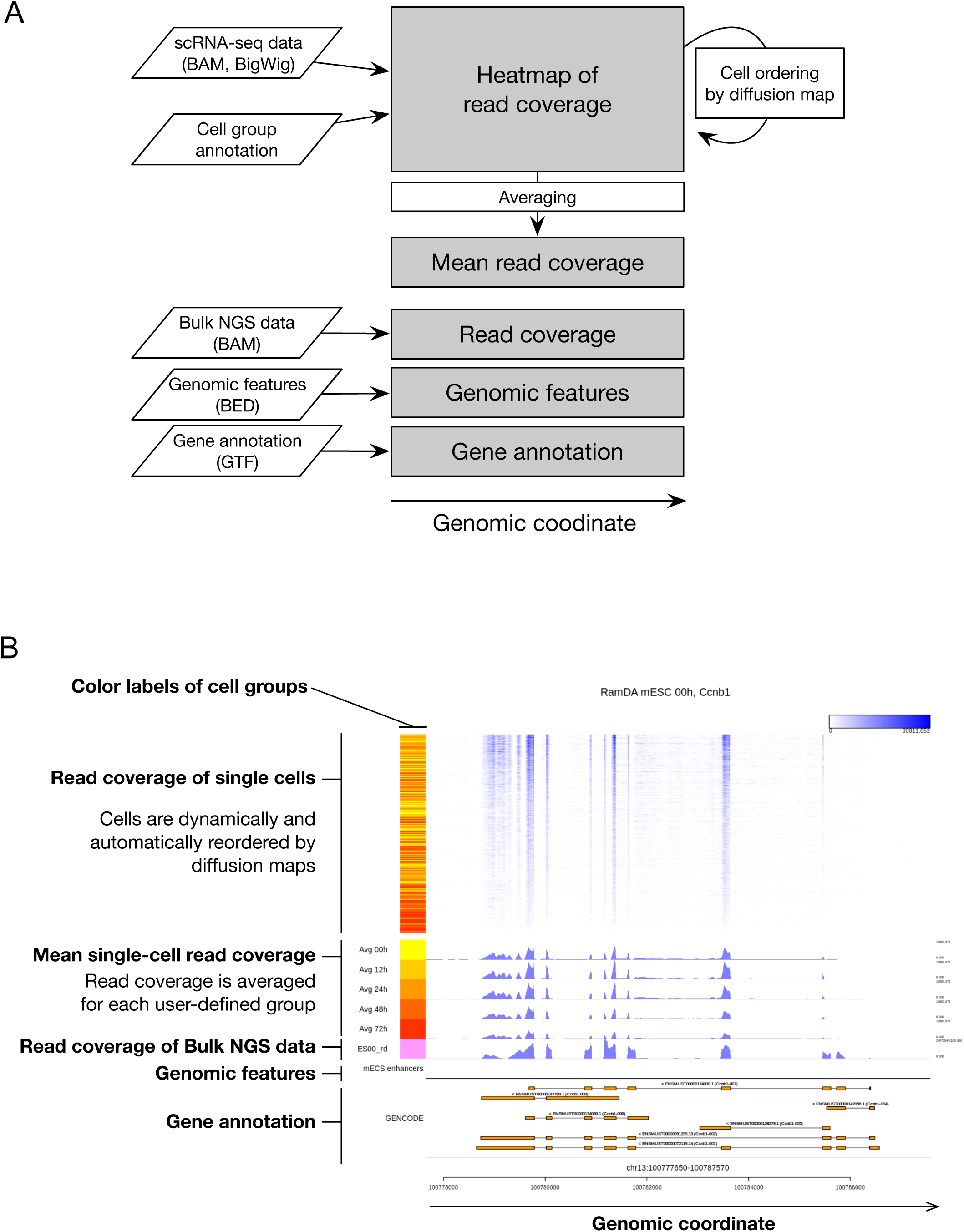
Overview of Millefy. (A) Millefy imports scRNA-seq data and visualizes read coverage of individual cells as a heat map. The rows (i.e., cells) of the heat map are dynamically reordered by diffusion maps. This automatic reordering highlights cell-to-cell heterogeneity in read coverage, which is hidden by mean read coverage data. Millefy associates genomic contexts, including bulk NGS data, genomic features, and gene annotations, thus facilitating the interpretation of single-cell read coverage. The parallelograms represent different types of input data. Gray boxes represent displayed tracks in Millefy. White boxes represent computation of a read coverage matrix. (B) An example plot of Millefy.

Millefy was implemented in R and can import scRNA-seq data without the need for format conversion. For scRNA-seq data, Millefy accepts BAM and BigWig formats, which are standard file formats for NGS data analysis. Millefy is dependent on the rtracklayer package [20] and Rsamtools package [21] for importing BAM and BigWig files, respectively. For gene annotation data and genomic features, Millefy accepts GTF and BED formats, respectively. The data.table package [22] is used to import GTF and BED files. For performing diffusion maps on read coverage data, Millefy utilizes the destiny package [23].

We provide Millefy as an R package and as a Docker image based on Jupyter Notebook Data Science stack (https://github.com/jupyter/docker-stacks) to facilitate installation and exploratory data science using Millefy in Jupyter notebooks.

## Results and Discussion

### Millefy highlights cellular heterogeneity in gene expression and transcribed gene structures

Researchers often merge read alignment files of single cells and visualize “synthetic bulk” data using standard genome browsers. However, in such cases, the merged (or averaged) read coverage cannot capture heterogeneity in read coverage. For example, a change in the merged read coverage cannot indicate whether the number of cells expressing a gene increased or the expression level of that gene increased across all cells. In contrast, Millefy visualizes read coverage of all individual cells in a scRNA-seq dataset as a heat map and thereby provides detailed information on cellular heterogeneity in read coverage.

To demonstrate the usefulness of Millefy’s ability to visualize read coverage in scRNA-seq data, we used a time-course RamDA-seq dataset derived from mouse embryonic stem cells (mESCs) upon induction of cell differentiation to primitive endoderm cells (at 0, 12, 24, 48, and 72 h) [12]. The dataset consists of 421 single cells.

Figure 2 shows the read coverage at *Sox17*, a differentiation marker gene. Cells were reordered according to the first diffusion component values calculated by a diffusion map of read coverage data for the locus, either within user-defined cell groups (Figure 2A) or across all cells (Figure 2B). While the height of the mean read coverage increased along the differentiation time course, the reordered heat map highlights the heterogeneity of read coverage among cells from the same time points (e.g., the 12 h group) (Figure 2). Specifically, Millefy showed that the number of cells with *Sox17* expression increased, indicating asynchronous cell differentiation progression among cells.

**Figure 2.**
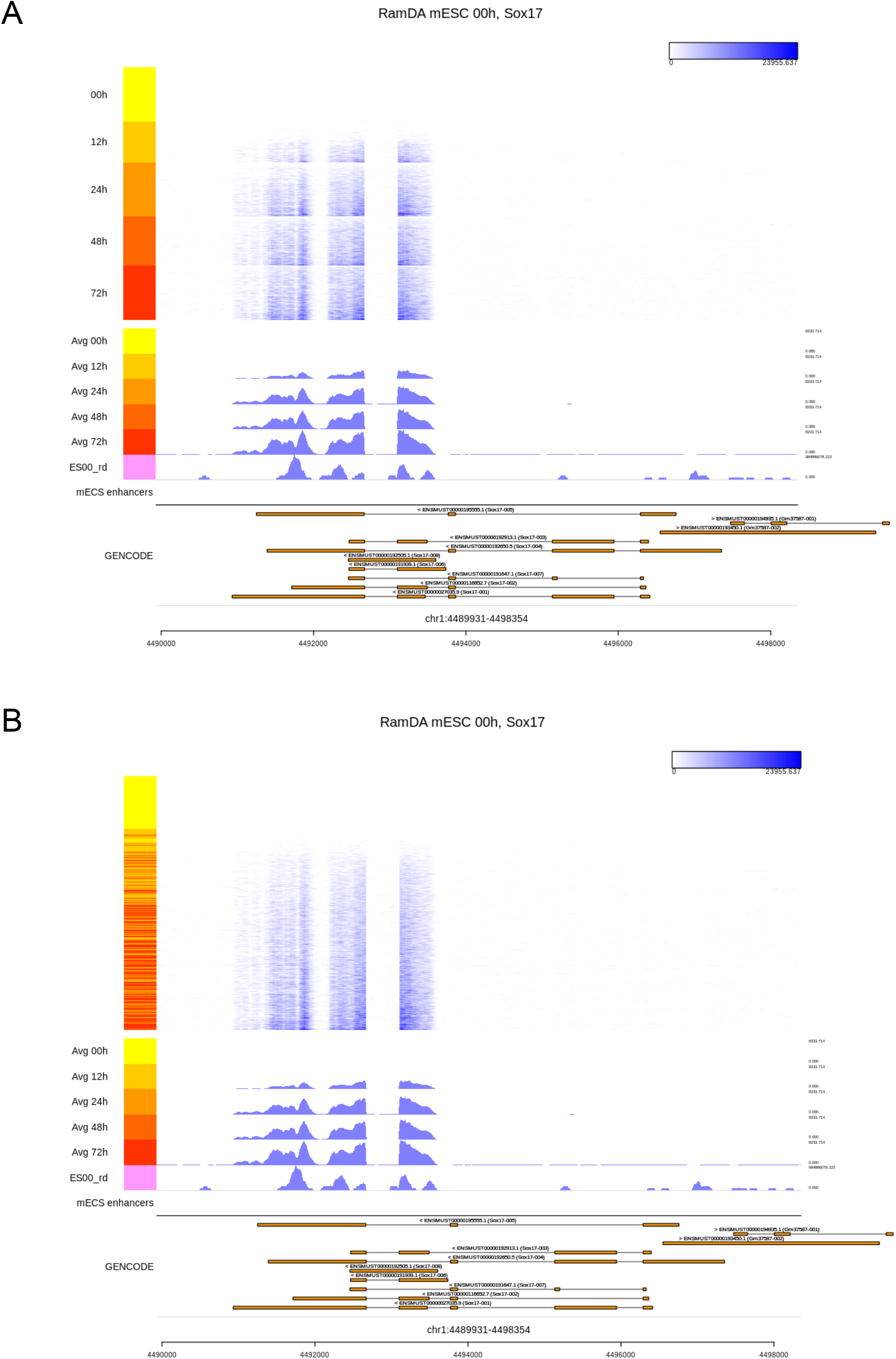
Millefy visualization of read coverage at the Sox17 locus. Millefy was applied to RamDA-seq data from mouse embryonic stem cells (mESCs) upon induction of cell differentiation to primitive endoderm cells. The top heat map shows single-cell read coverage. Color keys on the left side represent cells from different time points. The middle tracks show the averaged read coverage at different time points, the bulk RNA sequencing read coverage, and enhancer annotations. The bottom track shows the GENCODE reference gene annotation. Cells were reordered within (A) user-specified groups and (B) across all cells.

Another example is *Zmynd8*, a transcriptional repressor. Figure 3 shows read coverage of 421 individual cells at the *Zmynd8* locus. The cells were dynamically reordered using diffusion maps based on the read coverage in the focal region. Expression of the *Zmynd8* short isoform is known to be associated with the expression of its antisense RNA *Zmynd8as* [24]. While *Zmynd8as* is unannotated in the current gene annotation, the heat map by Millefy clearly showed differential regulation of the long isoform of *Zmynd8* and *Zmynd8as*, facilitating visual inspection of the two separated transcription units (Figure 3). We note that the averaged read coverage for each time point cannot distinguish whether the long and short isoforms of *Zmynd8* and *Zmynd8as* are correlated or uncorrelated. These results demonstrate that Millefy’s functionality for displaying read coverage as a reordered heat map reveals cell-to-cell heterogeneity at the focal locus.

**Figure 3.**
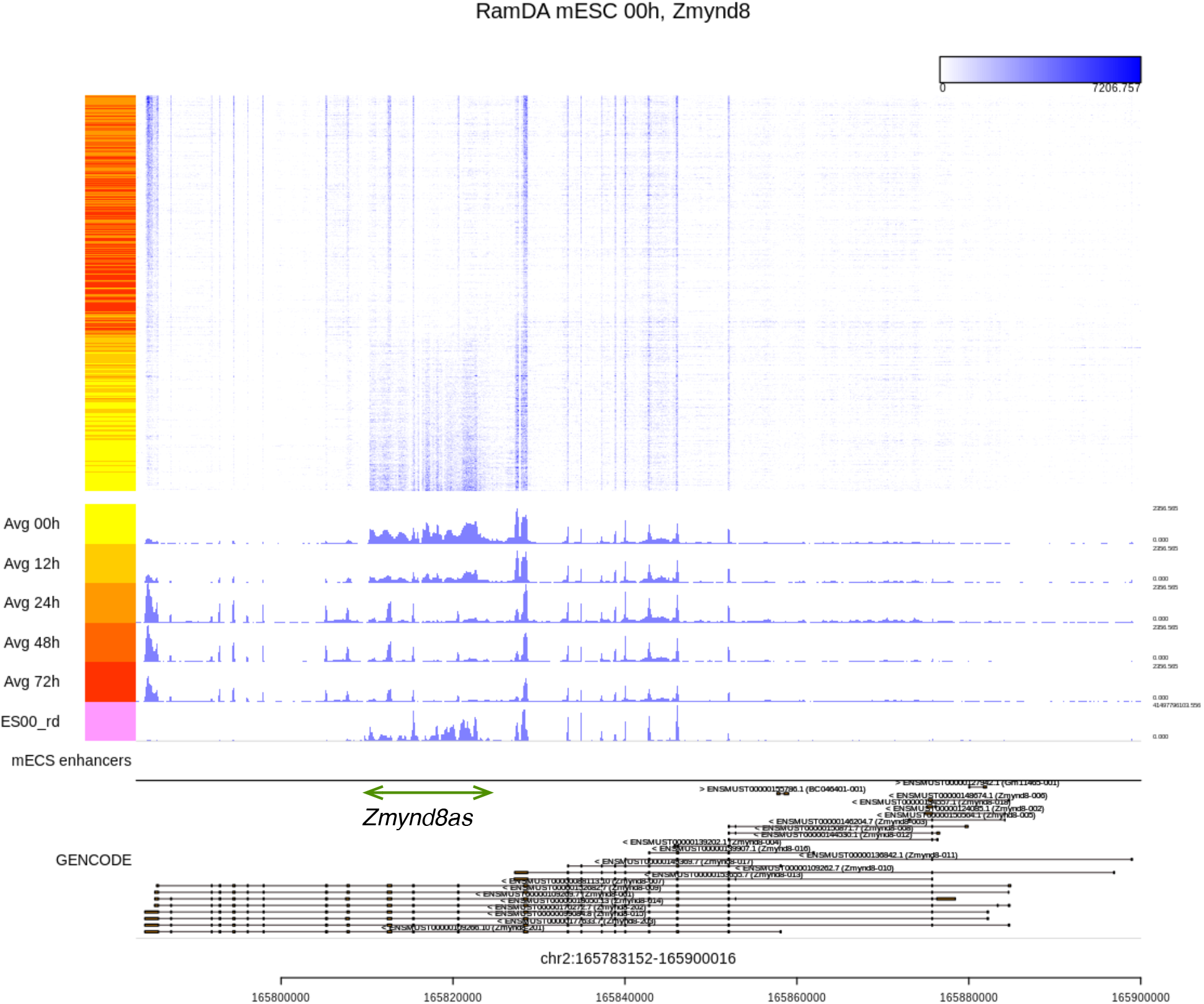
Visualization of the Zmynd8 locus by Millefy across all cells. Millefy was applied to RamDA-seq data from mouse embryonic stem cells (mESCs) upon induction of cell differentiation to primitive endoderm cells. All cells were reordered together without specified groupings. The green arrow has been added to indicate the position of *Zmynd8as*.

### Millefy associates read coverage with genomic contexts to facilitate interpretation of read coverage

Genomic contexts are crucial for interpretation of read coverage in bulk and single-cell RNA sequencing data. For example, read coverage overlapped with gene annotations can confirm known and reveal novel exon-intron structures. Moreover, read coverage overlapped with enhancer annotations can be interpreted as eRNA expression [6]. Using Millefy, single-cell read coverage can be compared with genomic and epigenomic features like enhancer elements.

To demonstrate the usefulness of the simultaneous visualization of single-cell read coverage and genomic contexts, we compared read coverage of the RamDA-seq data from mESCs (0 h) with mESC enhancer regions. We downloaded H3K4mel and H3K4me3 ChlP-seq peak regions for mESCs from the ENCODE project [25] and defined mESC enhancers as the H3K4mel peaks that (1) did not overlap with the H3K4me3 peaks, (2) were at least 2 kbp away from the transcriptional start sites, and (3) were outside of gene bodies of the GENCODE gene annotation (vM9) [26].

Figure 4 displays read coverage at the *Myc* locus, with the positions of enhancers active in mESCs. The *Myc* gene models and read coverage reveal that *Myc* was transcribed in mESCs. In addition, Millefy showed that some of the intergenic regions with transcribed RNA overlapped with the *Myc* downstream enhancer regions (Figure 4). This is consistent with the previous report that RamDA-seq can detect eRNAs [12]. This result exemplifies how Millefy can help to interpret read coverage of scRNA-seq data in genomic contexts.

**Figure 4.**
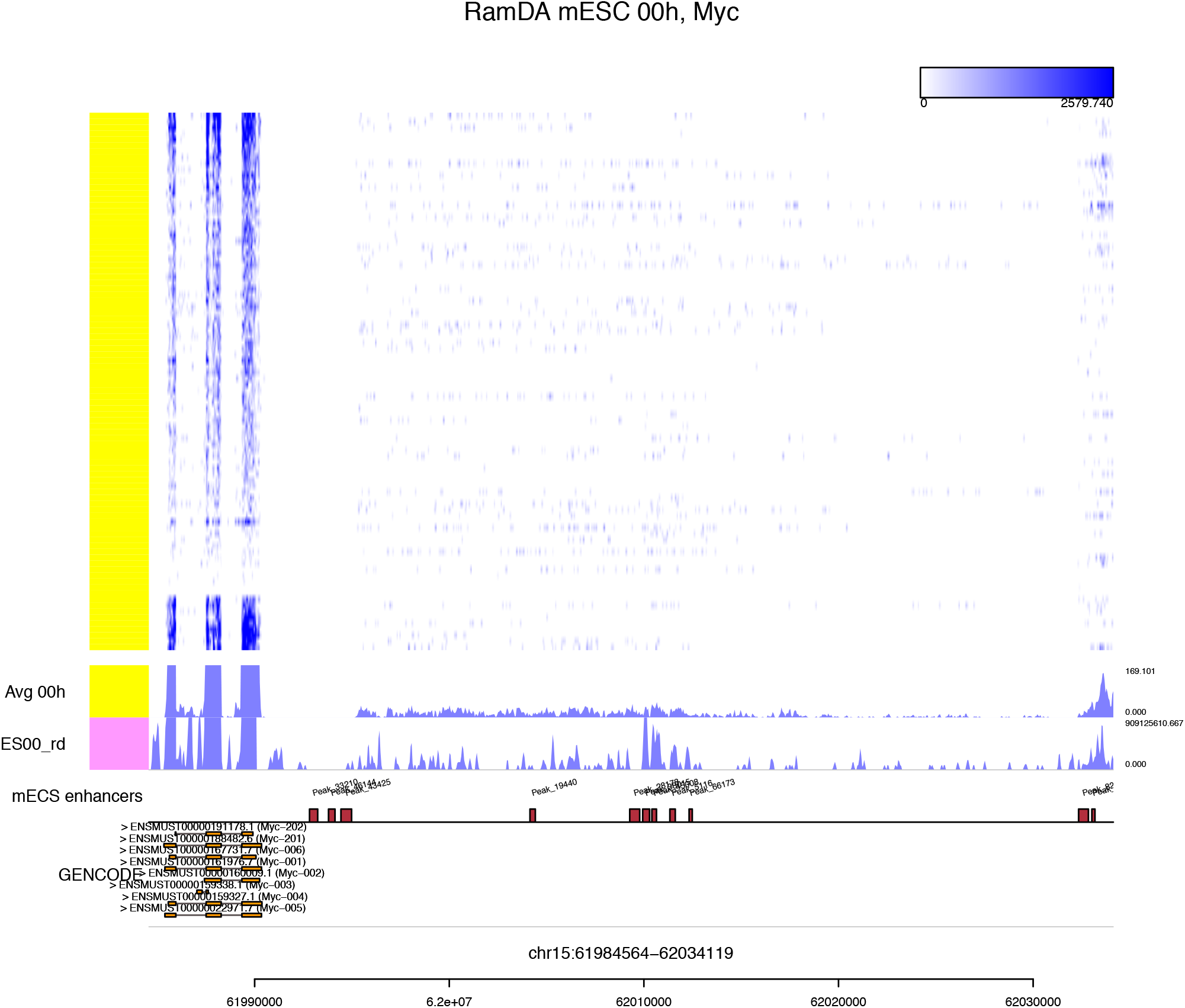
Millefy visualization of read coverage and enhancer regions around the Myc locus. Millefy was applied to RamDA-seq data from mouse embryonic stem cells (mESCs). All cells were reordered together without specified groupings. RNA transcription was observed for the enhancers on the left.

### Millefy facilitates quality control in full-length scRNA-seq methods

Millefy can also be used for QC in full-length scRNA-seq methods. For example, scRNA-seq read coverage of long transcripts indicates whether the method employed provided full-length transcript coverage. Full-length transcript coverage provides accurate information about isoform expression and gene structures and is a fundamental feature of full-length scRNA-seq methods [27].

We applied Millefy to C1-RamDA-seq data (*n* = 96) and C1-SMART-Seq V4 (*n* = 95) data from a dilution of 10 pg of mESC RNA. Figure 5 shows the read coverage at *Mdn1*, a gene with a long transcript (17,970 bp) consisting of 102 exons. C1-RamDA-seq detected all exons in most samples, while C1-SMART-seq V4 failed to detect a fraction of known exons. Interestingly, the patterns of missing exons in C1-SMART-seq V4 seemed to vary among the samples. The lower reproducibility in read coverage of C1-SMART-seq V4 relative to C1-RamDA-seq is likely owing to technical noise because the samples were prepared not from living cells but from a dilution of 10 pg of RNA. We note that mean read coverage cannot provide such detailed information on reproducibility in read coverage (Figure 5). This result demonstrates that Millefy visualizes read coverage in scRNA-seq as a QC measure and complements existing scRNA-seq QC pipelines based primarily on gene expression matrices [3].

**Figure 5.**
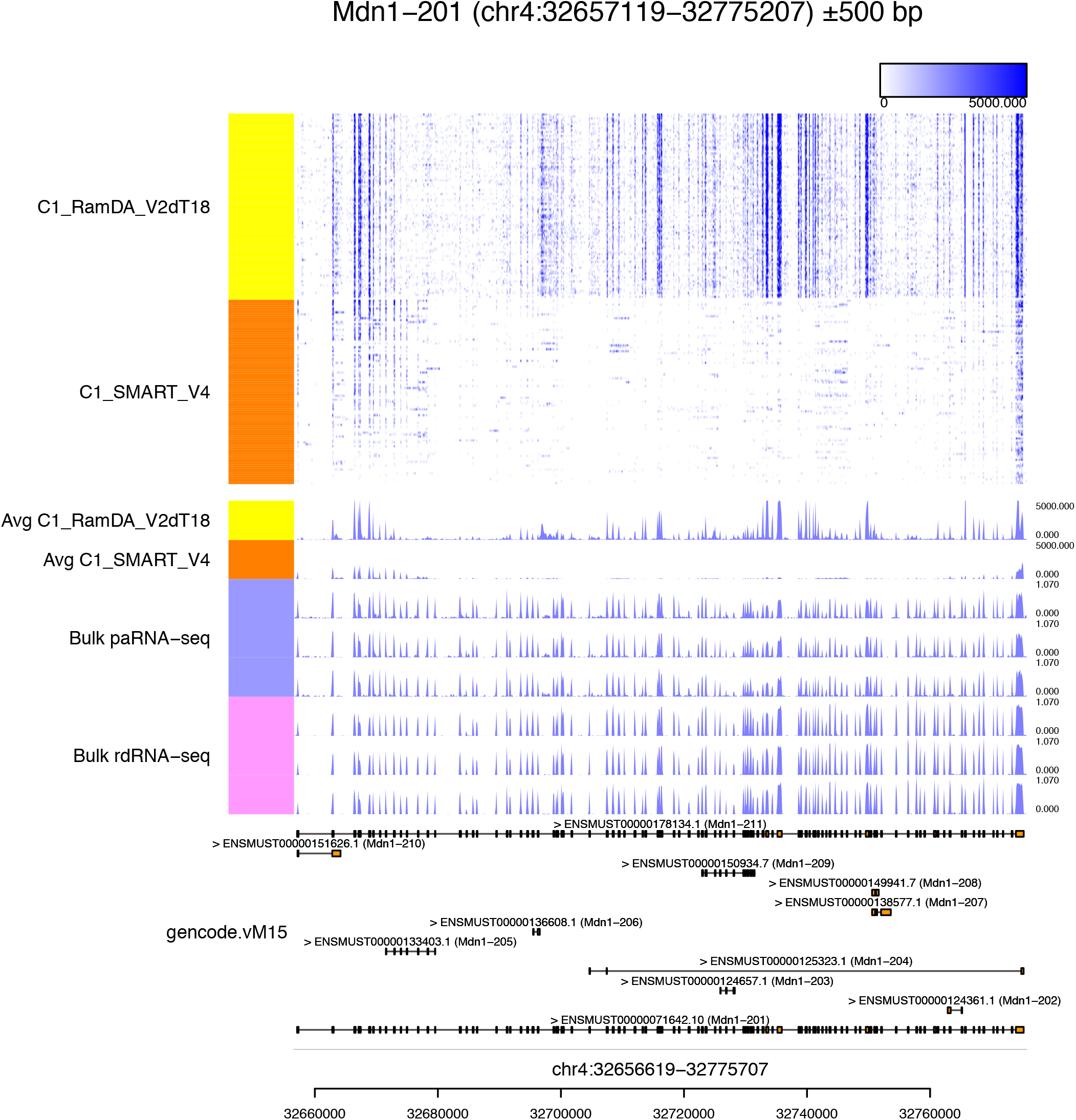
Example of quality control of scRNA-seq methods. Visualization of read coverage from C1-RamDA-seq (*n* = 95) and C1-SMART-Seq V4 (*n* = 96) data from a dilution of 10 pg of RNA at the *Mdn1* locus by Millefy. The samples were reordered within user-specified groups.

### Computational time

We measured the computational time of Millefy for visualizing whole gene bodies using RamDA-seq data with 793 samples from mESCs [12]. For 1000 randomly selected gene loci (of expressed genes with average TPM>5), Millefy processed BigWig and BAM files in 39.3 and 138.9 s, respectively, on average.

## Conclusions

Millefy, which is integrated with Jupyter Notebook and provided as a Docker image, can easily be utilized in exploratory analyses of scRNA-seq data. Moreover, in the development of bioinformatics methods using rule-based and machine learning approaches for profiling alternative splicing or novel RNAs by scRNA-seq, visualization of read coverage will become more important for evaluating and representing the predictions of algorithms. In conclusion, Millefy can help researchers assess cellular heterogeneity and RNA biology using scRNA-seq data.

## Availability and requirements

**Project name**: Millefy

**Project home page**: https://github.com/yuifu/millefy (R package), https://hub.docker.com/r/yuifu/datascience-notebook-millefy (Docker image)

**Archived version**: DOI:10.5281/zenodo.2555019

**Operating system(s)**: Platform independent

**Programming language**: R

**Other requirements**: R version 3.2.2 or higher

**License**: MIT

**Any restrictions to use by non-academics**: No

## List of abbreviations

eRNAs: enhancer RNAs
mESCs: mouse embryonic stem cells
PCA: principal component analysis
QC: quality control
scRNA-seq: single-cell RNA sequencing

## Ethics approval and consent to participate

Not applicable.

## Consent for publication

Not applicable.

## Availability of data and materials

The datasets supporting the conclusions of this article are available in the Gene Expression Omnibus repository, GSE98664.

## Authors’ contributions

HO designed and implemented the software, performed the data analyses, and wrote the manuscript. TH and MU performed the C1-RamDA-seq and C1-SMART-Seq V4 experiments. IN contributed to the design of the data analyses. All authors read and approved the final manuscript.

## Acknowledgements

We thank the members of the Bioinformatics Research Unit, particularly Hirotaka Matsumoto and Mika Yoshimura, for discussion of data analyses and Manabu Ishii and Akihiro Matsushima for IT infrastructure management.

